# QuPath: Open source software for digital pathology image analysis

**DOI:** 10.1101/099796

**Authors:** Peter Bankhead, Maurice B Loughrey, José A Fernández, Yvonne Dombrowski, Darragh G McArt, Philip D Dunne, Stephen McQuaid, Ronan T Gray, Liam J Murray, Helen G Coleman, Jacqueline A James, Manuel Salto-Tellez, Peter W Hamilton

## Abstract

QuPath is new bioimage analysis software designed to meet the growing need for a user-friendly, extensible, open-source solution for digital pathology and whole slide image analysis. In addition to offering a comprehensive panel of tumor identification and high-throughput biomarker evaluation tools, QuPath provides researchers with powerful batch-processing and scripting functionality, and an extensible platform with which to develop and share new algorithms to analyze complex tissue images. Furthermore, QuPath’s flexible design makes it suitable for a wide range of additional image analysis applications across biomedical research.

## Introduction

The ability to acquire high resolution digital scans of entire microscopic slides with high-resolution whole slide scanners is transforming tissue biomarker and companion diagnostic discovery through digital image analytics, automation, quantitation and objective screening of tissue samples. This area has become widely known as digital pathology^1,2^. Whole slide scanners can rapidly generate ultra-large 2D images or z-stacks in which each plane may contain up to 40 GB uncompressed data. Manual subjective scoring of this data by traditional pathologist assessment is no longer sufficient to support large-scale tissue biomarker trials, and cannot ensure the high quality, reproducible, objective analysis essential for reliable clinical correlation and candidate biomarker selection. New and powerful software tools are urgently required to ensure that pathological assessment of tissue is practical, accessible and reliable for biological discovery and the development of clinically-relevant tissue diagnostics.

In recent years, a vibrant ecosystem of open source bioimage analysis software has developed. Led by ImageJ^3^, researchers in multiple disciplines can now choose from a selection of powerful tools, such as Fiji^4^, Icy^5^, and CellProfiler^6^, to perform their image analyses. These open source packages encourage users to engage in further development and sharing of customized analysis solutions in the form of plugins, scripts, pipelines or workflows - enhancing the quality and reproducibility of research, particularly in the fields of microscopy and high content imaging. This template for open-source development of software has provided opportunities for the field of image analysis to add considerably to translational research by enabling the development of the bespoke analytical methods that are essential to address specific issues. There has, however, been an absence of tools to tackle the specific visualization and computational challenges posed by whole slide images (WSI) and very large 2D data, and this has created a major bottleneck in the pathological tissue assessment pipeline^7^. Consequently, the field of digital pathology continues to lack a commonly-accepted, open and accessible software framework for distributing novel algorithms, and has been slow to embrace the ‘call for bioimaging software usability’^8^. In practice, this has meant that users without access to expensive commercial solutions have had to either resort to inefficient workarounds (such as image downsampling and cropping) to be able to apply general open source analysis tools to a subset of their data^9,10^, or to rely primarily on laborious manual evaluation of slides, which is known to have high variability and limited reproducibility^11,12^.

QuPath (https://qupath.github.io) has been developed to address these needs by offering the first comprehensive, open source platform specifically designed for whole slide applications. At its core is a cross-platform, multithreaded, tile-based whole slide image viewer, which incorporates extensive annotation and visualization tools. On top of this, QuPath offers an array of novel algorithms to provide not only ready-made, user-friendly solutions to common, challenging analysis problems in pathology, but also the building blocks to create custom workflows - and link these together for batch processing with powerful scripting functionality **(Fig. 1)**. Finally, QuPath enables developers to add their own extensions to solve new challenges and applications, and to exchange data in a streamlined manner with existing tools that otherwise lack whole slide support, such as ImageJ and MATLAB.

**Figure 1.**
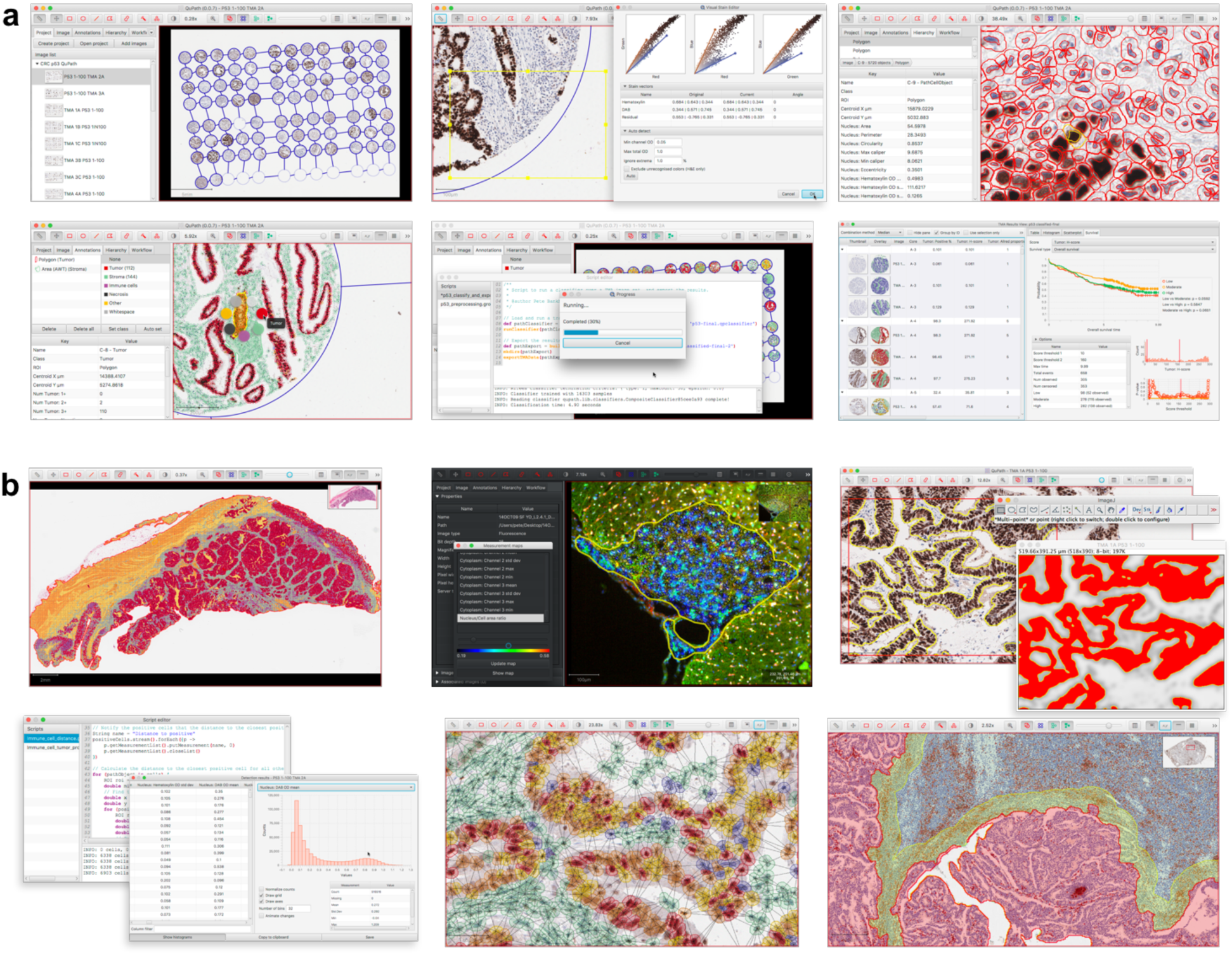
Illustration of QuPath’s use and functionality. (**a**) A typical workflow for TMA analysis (here, p53) demonstrates several of QuPath’s main features (left-to-right): Creation of a multi-slide project with automated TMA dearraying, stain estimation, cell detection and feature computation, trainable cell classification, batch processing, and survival analysis. (**b**) QuPath offers a wide range of additional functionality, including support for whole face tissue sections and fluorescence image analysis, data exchange with existing software and platforms (e.g. ImageJ and MATLAB), scriptable data mining, and rapid generation, visualization and export of spatial, morphological and intensity-based features.

A key feature underpinning QuPath’s functionality, and a major technical distinguishing factor between QuPath and other bioimaging analysis software, is its hierarchical, ‘object-based’ data model. Here, an ‘object’ refers to a structure or region within the image, which may be created and manipulated by either interactive drawing tools (e.g. to annotate a particular region of interest) or automated segmentation commands (e.g. to detect individual nuclei or cells). This generic model allows QuPath to represent and display relationships between very large numbers of image objects in an efficient and intuitive manner across gigapixel images, and support the fast and interactive training of object classifiers using machine learning techniques.

A practical example of this is in the evaluation of the presence, localization and intensity of expression of key diagnostic, prognostic and predictive biomarkers in tissue sections. These biomarkers are typically detected using antibodies and chromogenic based detection systems, and are selectively expressed in tumor cells or in other cellular compartments. QuPath’s built-in cell segmentation algorithms can detect potentially millions of cells as objects within a single WSI, in addition to measuring cell morphology and biomarker expression. QuPath further supports the classification of different cell types according to these features, to generate a comprehensive phenotypic description of each cell within the tissue sample. This in turn provides a quantitative cellular map of the entire tissue section, which can be subsequently selected, queried and filtered to mine the image data and uncover morphological subtleties not immediately visible during traditional pathological assessment. All of this can typically be achieved within minutes, without a requirement for specialist hardware.

## Results and Discussion

To demonstrate some of these capabilities, including its biological and clinical validity, we used QuPath to analyze several image sets derived from surgical resection specimens from a population-based cohort of 660 patients with stage II and stage III colon cancer, diagnosed between 2004-2008 (392 stage II, 268 stage III) and with high-quality curation of clinicopathological information. From representative paraffin-embedded tumor blocks, provided via the Northern Ireland Biobank, whole sections were haematoxylin and eosin-stained in the Northern Ireland Molecular Pathology Laboratory. Tissue microarrays (TMAs) were generated from representative 1mm diameter tumor cores from each case, sampled in triplicate from the tumor center after annotation.

The aim was to test the performance of QuPath in the context of known and novel biological and clinical data, using cancer immunology and tumor suppressor genes as paradigms. Firstly, we applied QuPath to analyze TMAs immunohistochemically-stained (IHC) for the T cell markers CD3 and CD8. Following digital scanning of the WSI, the initial QuPath setup required importing the images, applying automated ‘dearraying’ to identify tissue cores, manually refining the resulting dearrayed grid, and removing cores unsuitable for analysis. After this step, batch analysis was applied across all TMA slides to identify the tissue within each core and automatically count the number of positive cells per mm^2^ tissue based upon a fast peak-finding algorithm after stain separation by color deconvolution^13^ (**Supplementary Fig. 1, Supplementary Video 1**). Running on a standard Mac Pro (3.5 GHz, 6-Core Intel Xeon E5, 32-GB RAM), this approach required less than 4 minutes for each biomarker to analyze 21 whole slide images, representing a total of > 2000 tissue cores per biomarker and counting approximately 1.2 million CD3 positive and 0.6 million CD8 positive cells, while simultaneously exporting low-resolution images of each core both with and without markup for verification. After applying a median cutoff to the exported results, a statistically significant association between disease-specific survival and positive cell density scores was demonstrated for both CD3 and CD8 (log-rank test, p-values 0.006 and 0.007 respectively; **Fig. 2a-b**). This recapitulates within our cohort the seminal work of Galon *et al* in demonstrating the prognostic relevance of adaptive immunity in colorectal cancer^14^, while demonstrating a highly-efficient method of investigating similar markers within other cohorts and cancer types.

**Figure 2.**
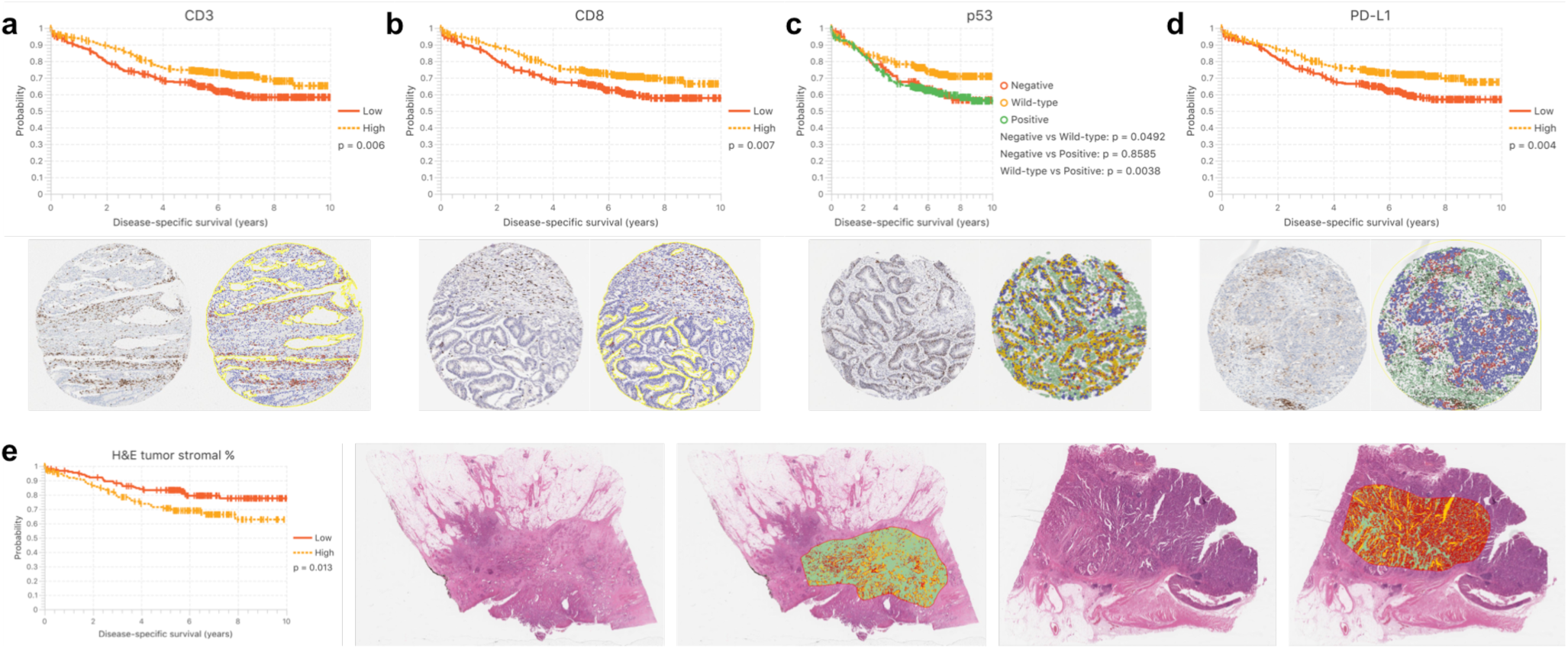
Survival analysis of colon cancer cohort based on QuPath automated image analysis. (**a-d**) Kaplan Meier survival analysis for biomarker scores of TMAs stained for CD3, CD8, p53 and PD-L1. Median cutoffs are applied in all cases, except p53 where two cutoffs were selected by an experienced pathologist to distinguish between aberrant negative, “wild type” (normal) and aberrant positive groups. Representative images showing an original core and QuPath markup image are included below. (**e**) Kaplan Meier curve showing patient stratification based on median tumor stromal percentage. Representative images show the original images and markup for tumors with a high and low stromal percentage respectively. Green indicates regions classified as stroma, dark red indicates tumor epithelium, while yellow represents other classified tissue or whitespace.

Next, we used QuPath to evaluate immunohistochemistry for p53 in a second set of TMAs from the same cohort. This required a more sophisticated analysis to encompass the biological understanding and staining pattern of the marker. After applying QuPath’s cell detection algorithm to segment and measure cells within each core, a random trees classifier^15^ was interactively trained to enable p53 expression to be scored selectively within the epithelial cell population according to nuclear staining intensity and proportion (**Supplementary Fig. 2, Supplementary Video 2**). This analysis showed that aberrant p53 expression (diffuse intense or completely absent immunoreactivity) is associated with significantly poorer unadjusted disease-specific survival when compared with intermediate, ‘wild-type’ expression (p=0.003 extreme negative/positive vs. intermediate; **Fig. 2c**). Despite the well-established role of TP53 in colorectal cancer carcinogenesis, results from prognostic studies assessing p53 IHC expression have been inconsistent^16^. However, the extreme negative pattern of aberrant p53 immunoreactivity has only been described relatively recently^17,18^ and has not been widely assessed in colorectal cancer cohorts. This example therefore emphasises the flexibility of the QuPath open source platform in measuring common IHC markers with variable tumor expression patterns, and demonstrates how the relationship between quantitative cellular analysis and clinical outcome can be robustly assessed.

We then applied QuPath to the analysis of programmed cell death ligand 1 (PD-L1) immunoexpression in the same TMA cohort. PD-L1 immunoexpression is a prognostic marker in a range of cancer types and also a predictor of response to immune checkpoint therapy in some cancers^19,20^. However, there is a lack of consensus on the epitopes of clinical relevance and, more importantly, the optimal scoring systems for evaluation. The approach to analysis was similar in principle to that adopted for p53, however further attention was required because of additional challenges posed by PD-L1 immunostaining. Firstly, staining is cytoplasmic and/or membranous rather than nuclear. Secondly, PD-L1 can be expressed in tumor epithelium, but is more commonly expressed in other tissue compartments, notably within peritumoral stromal inflammatory cells. Although this level of heterogeneity in staining pattern is increasingly being recognised, the clinical importance of distinguishing tumor epithelial from inflammatory cell staining is yet to be fully understood. Sufficient cell classification is therefore required to identify both positively and negatively staining tumor (epithelial) and non-tumor cell populations, in addition to distinguishing true protein expression levels from the various staining artefacts that are inherent with IHC-based tissue analysis. Here, applying QuPath, a cell was classified as positive or negative based on maximal DAB staining intensity, as a surrogate marker of protein expression, within a full cell region approximated by expanding detected nuclei (**Supplementary Fig. 3**). Across the cohort, a median cutoff of 1.46% of all cells exhibiting PD-L1 positivity according to this definition was applied to stratify patients, showing high PD-L1 expression to be significantly associated with improved disease-specific survival in unadjusted analysis (p=0.004, **Fig. 2d**). Additionally, an analysis based on tertiles exhibited a dose-response effect, and separation of the tumor (epithelial) and non-epithelial components suggested that PD-L1 expression in colon cancer tissue is primarily found in the non-epithelial compartment (**Supplementary Fig. 4**). These results support the incipient evidence of PD-L1 prognostic value in colorectal cancer reported by ourselves and others in independent cohorts^20–22^, and may be of help when used together with tumor microsatellite instability status for patient stratification in consideration of anti-PD-Ll therapy.

Finally, to demonstrate QuPath’s flexibility beyond IHC scoring in TMAs, we applied texture-based analysis to calculate the tumor stromal percentage in whole face representative tumor sections stained with hematoxylin and eosin (H&E) from 312 patients with stage II colon cancer from the same cohort. Several studies have reported that a higher intratumoral stromal percentage correlates with a worse prognosis in patients with stage II and stage III colorectal cancer^23,24^. Given highly variable clinical outcomes within stage II disease, there is a particular need for additional prognostic features in this group, to identify those patients most likely to benefit from adjuvant chemotherapy. These studies employed a crude visual estimate of tumor stromal percentage. Applying QuPath’s more fine-grained and reproducible assessment, and a median cutoff for analysis, we found lower disease-specific survival among patients with a high tumor stromal percentage in unadjusted analysis (p=0.013; **Fig. 2e**), consistent with previous findings^23,24^.

In summary, QuPath is the first comprehensive, open source bioimage analysis platform designed for whole slide images. We have demonstrated that it provides the tools necessary for fast, accurate and reproducible digital pathology analysis across a range of challenging applications. All of the above represent analyses that are cumbersome and time-consuming for pathologists to perform manually, given that they depend upon the accurate visual estimation of proportional staining within large numbers of stained cells, or of proportional composition of complex tissue areas. Accurate assessment is particularly difficult when the clinically relevant cutoff is very low and expression very localized (e.g. PD-L1), requiring detailed examination and precision. Such precise analyses are becoming increasingly necessary and important, both clinically in the era of personalized medicine, and in a research context for high-throughput evaluation of novel biomarkers. By offering an extensible environment for pathologists, biologists, and computer scientists to build highly performant algorithms for image interpretation and analysis, there is potential to drive adoption of quantitative imaging in academic, diagnostic and pharmaceutical research organizations, and to accelerate biomarker discovery in large scale multinational clinical trials. This is an absolute requirement to ensure a return on the global investment made in companion diagnostics for precision medicine.

## Materials and Methods

### Cohort

This analysis was based on a population-based cohort of 660 stage II/III colon adenocarcinoma patients, representing 89% of all patients undergoing surgery for stage II/III colon adenocarcinoma in two Healthcare Trusts in Northern Ireland between 2004 and 2008. Patients were sampled and clinical information obtained from the Northern Ireland Cancer Registry. All patients were followed up for occurrence and cause of death via linkage to the Northern Ireland Registrar General’s Office up to 31^st^ December 2013. Ethical approval for these linkages was given by ORECNI (REC: 10/NIR02/53). Over ten years (and a mean of 5 years) of clinical follow-up, 46% of patients had died, of which 38% were from CRC-specific causes. Corresponding tumor slides and blocks were retrieved and collated via the Northern Ireland Biobank (REC:11/NI/0013).

Following slide review, a new section was cut for H&E staining from a single representative tumor block in each case, and the new slides annotated for tissue microarray (TMA) construction. Three representative areas within the tumor center of each block were annotated for targeted coring (by an experienced biomedical scientist and confirmed by expert pathologists, MBL and JAJ). One millimeter diameter tissue cores were extracted from donor blocks and inserted into recipient blocks using a manual tissue arrayer (Estigen, Tartu, Estonia).

### Immunohistochemistry

All IHC was performed in a hybrid laboratory (Northern Ireland Molecular Pathology Laboratory) that has UK Clinical Pathology Accreditation. Internally validated biomarker conditions, which followed UK-NEQAS guidelines (CD3 and CD8) or were based on expected performance from the literature (p53 and PD-L1), were as follows: CD3 (clone 2GV6 Ventana BenchMark; CC1 32minutes, Optiview detection), CD8 (clone c8/144B, Dako: Leica Bond III, ER2 20mins, 1/50, polymer detection), p53 (clone D0-7, Dako, ER2 30 mins, 1/100, polymer detection), PD-L1 (clone SP142, Ventana BenchMark, CC1 24mins, optiview detection).

### Image data acquisition

All TMA slides were scanned using an Aperio ScanScope CS whole slide scanner at 40X magnification, with a resolution of 0.25 μm/pixel. H&E slide scanning was heterogenous: 231 were scanned on the Aperio ScanScope scanner, while 81 were scanned on a Hamamatsu Nanozoomer. The resolution of all images was within the range 0.231 – 0.253 μm/pixel.

### Software

QuPath was written as a novel, cross-platform Java application. The core software was developed using Java 8, with a user interface written using JavaFX. Whole slide image reading was achieved using QuPath’s interfaces to the OpenSlide library^25^, while the implementations of QuPath’s cell detection and superpixel segmentation commands made use of ImageJ as a library for standard image processing operations^3^. The random trees classifier and fast cell counting were implemented using the OpenCV library (http://opencv.org). The analysis for this study was performed on a Mac Pro (3.5 GHz, 6-Core Intel Xeon E5, 32-GB RAM).

### Code availability

Source code and documentation for QuPath are available at https://qupath.github.io

### Statistical analysis

Survival curves using the Kaplan Meier method were generated and log-rank tests applied using the *TMA data viewer* within QuPath, and independently verified using R (version 3.2.2)^26^ with the ‘Survival’ package (version 2.38-3)^27^. For the calculation of disease-specific survival, deaths from other causes were treated as censored events. Median cutoff values were used in all cases, except for p53 where an experienced pathologist (MBL) selected two biologically-plausible cutoffs (H-scores 10 and 160) to separate extreme positive and extreme negative cases from those with intermediate (‘wild-type’) expression, based upon viewing all TMA cores post-analysis ranked by H-score. Stratification based on tertiles is also provided in the Supplementary Materials for PD-L1.

For TMA analysis, up to three tissue cores were available from each tumor, all selected from the same paraffin block representing the central tumor region. A single patient biomarker score was defined as the median of all available scores for the corresponding patient and biomarker. The median was chosen to aid the robustness of the measurement in a high-throughput setting, and reduce the likelihood of basing the score for any individual patient on an outlier that may have been caused by a tissue or staining artefact.

### Tissue microarray preprocessing

Separate projects were created within QuPath for each biomarker, and the slide images imported to the corresponding projects. QuPath’s automated *TMA dearrayer* was applied in batch over all slides within each project to identify tissue cores. The resulting TMA grid was manually verified and amended where necessary, e.g. to adjust the locations of cores that were outside their expected position, or to remove cores where prominent artefacts were visible. Patient identifiers were then imported into QuPath for each core to assist alignment with survival data later. Additionally, stain vector (i.e. color) and background estimates were applied for each IHC analysis project to improve stain separation within QuPath using color deconvolution^13^. This was achieved by selecting a representative area containing an area of background along with examples of strong hematoxylin and DAB staining, and applying QuPath’s *Estimate stain vectors* command to identify stain vectors within this region. The resulting vectors were then used for all images in the project.

### Analysis of CD3 and CD8 IHC

After the initial TMA preprocessing steps described above, analysis of CD3 and CD8 was performed using QuPath’s *Simple tissue detection* and *Fast cell counts* commands. Briefly, tissue was detected within each TMA core by thresholding a downsampled and smoothed image of the core, followed by cleanup of the resulting binary image by morphological operations. Individual cells were identified by separating stains using color deconvolution and identifying peaks in either the hematoxylin channel (CD3) or sum of the hematoxylin and DAB channels (CD8) after smoothing, and assigning these as positive or negative cells based upon the smoothed DAB channel information. The number of positive cells and area detected were used to calculate the average number of positive cells per mm^2^, and these results exported along with markup images showing the detected cells, for visual verification. The detection and export steps were fully automated using a batch processing script.

### Analysis of p53 IHC

Preprocessing steps were applied as described above. QuPath’s *Cell detection* command was then used to identify cells across all cores based upon nuclear staining. This command additionally estimates the full extent of each cell based upon a constrained expansion of the nucleus region, and calculates up to 33 measurements of intensity and morphology, including nucleus area, circularity, staining intensity for hematoxylin and DAB, and nucleus/cell area ratio. A subset of these measurements was chosen and supplemented for each cell by measuring the local density of cells, and taking a Gaussian-weighted sum of the corresponding measurements within neighboring cells using QuPath’s *Add smoothed features* command. A two-way random trees classifier was then interactively trained to distinguish tumor epithelial cells from all other detections (comprising non-epithelial cells, necrosis, or any artefacts misidentified as cells) and applied across all slides (see **Supplementary Video 2**). Intensity thresholds were set to further subclassify tumor cells as being negative, weak, moderate or strongly positive for p53 staining based upon mean nuclear DAB optical densities. An H-score was calculated for each tissue core by adding 3x% strongly stained tumor nuclei, 2×% moderately stained tumor nuclei, and 1× % weakly stained tumor nuclei^28^, giving results in the range 0 (all tumor nuclei negative) to 300 (all tumor nuclei strongly positive).

### Analysis of PD-L1 IHC

The approach to scoring PD-L1 was similar to p53, apart from the following: 1) a three-way random trees classifier was trained to distinguish between epithelial, non-epithelial and ‘other’ detections (including artefacts and necrosis); 2) cells were classified as positive or negative based upon a single intensity threshold applied to the maximum DAB optical density within the full cell area, and 3) summary scores were generated as the percentage of cells classified as positive, with ‘other’ detections removed.

### Analysis of tumor stromal percentage in H&E whole face sections

Representative tumor regions were annotated across all 312 H&E-stained slides by an experienced pathologist (MBL) using QuPath’s manual annotation tools. A script was then applied in batch to automatically identify and set the average background intensity for the red, green and blue channels of each image, which varied markedly according to the scanner used. A second script was then run over all images to apply QuPath’s *SLIC superpixel segmentation* command to subdivide each annotated region into ‘superpixels’ based upon simple linear iterative clustering^29^. This script additionally calculated both the average hue for each superpixel along with Haralick texture features^30^ from optical density values using QuPath’s *Add intensity features* command. QuPath’s *Add smoothed features* command was also applied to calculate a Gaussian-weighted sum of the features of neighboring superpixels, and append these to the existing features for each superpixel. This provide additional contextual information extending beyond the superpixel itself.

A subset of 40 ‘training’ images was then identified for the pathologist to interactively train a random trees classifier to distinguish between tissue areas comprising tumor epithelium, stroma and ‘other’ (e.g. whitespace, mucin, normal muscle or necrosis). This required drawing around regions containing tissue of each class and annotating these accordingly. During this process, QuPath used all available features to train the classifier in a background process and thereby provide immediate feedback on classification performance. Once the classification was considered adequate across the training images, the classifier was applied to all images within the set and the total area of superpixels for each class was exported. The tumor stromal percentage (TSP) was then calculated as

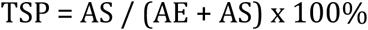

where AS represents the total area classified as stroma, and AE represents the total area classified as epithelium.

## Acknowledgements

The research leading to these results has received funding from Invest Northern Ireland (RDO0712612 to P.W.H.); Cancer Research UK Accelerator (C11512/A20256 to P.W.H./M.S.-T./J.A.J). The Northern Ireland Molecular Pathology Laboratory is supported by Cancer Research UK, Experimental Cancer Medicine Centre Network, the NI Health and Social Care Research and Development Division, the Sean Crummey Memorial Fund, the Tom Simms Memorial Fund and the Friends of the Cancer Centre (to M.S.-T.). The Northern Ireland Biobank is funded by the Health and Social Care Research and Development Division of the Public Health Agency in Northern Ireland and Cancer Research UK (to J.A.J/P.W.H/S.McQ ref. SPI/5069/14; additional support was received from the Friends of the Cancer Centre (to J.A.J./P.W.H/M.B.L). The cohort was supported by Cancer Research UK (ref. C37703/A15333 and ref. C50104/A17592) and the HSC R&D office of Northern Ireland (ref. EAT/4905/13). The Northern Ireland Cancer Registry is funded by the NI Public Health Agency.

We thank Mr Ken Arthur for the construction of the original tissue microarrays used in this study, Ms Victoria Bingham for her work in staining and scanning the slides, and both Dr Roisin O’Neill and the Northern Ireland Cancer Registry for their contributions to the clinical data collation.

## Author contributions

PB designed and developed the software. Feedback and testing of the software was provided by JAF, YD, DGM, PWH, PDD, and MBL. Before scanning and image analysis the expected expression of the biomarkers on tissue sections was initially assessed microscopically as fit-for-purpose by a team of experienced histopathologists MS-T, SMcQ, JAJ. Experiments were designed by PB, MBL, MS-T and PWH, and performed by PB and MBL. Data was analyzed by PB, JAF and HGC. Manuscript was written by PB with contributions from all authors.

## References

1. Pantanowitz, L. et al. Review of the current state of whole slide imaging in pathology. J. Pathol. Inform. 2, 36 (2011).

2. Hamilton, P. W. et al. Digital pathology and image analysis in tissue biomarker research. Methods 70, 59–73 (2014).

3. Schneider, C. a, Rasband, W. S. & Eliceiri, K. W. NIH Image to ImageJ: 25 years of image analysis. Nat. Methods 9, 671–675 (2012).

4. Schindelin, J. et al. Fiji: an open-source platform for biological-image analysis. Nat. Methods 9, 676–82 (2012).

5. de Chaumont, F. et al. Icy: an open bioimage informatics platform for extended reproducible research. Nat. Methods 9, 690–6 (2012).

6. Lamprecht, M., Sabatini, D. & Carpenter, A. CellProfiler™: free, versatile software for automated biological image analysis. Biotechniques 42, 71–75 (2007).

7. Heindl, A., Nawaz, S. & Yuan, Y. Mapping spatial heterogeneity in the tumor microenvironment: a new era for digital pathology. Lab. Investig. 95, 377–384 (2015).

8. Carpenter, A. E., Kamentsky, L. & Eliceiri, K. W. A call for bioimaging software usability. Nat. Methods 9, 666–70 (2012).

9. Deroulers, C. et al. Analyzing huge pathology images with open source software. Diagn. Pathol. 8, 92 (2013).

10. Nelissen, B. G. L., van Herwaarden, J. a., Moll, F. L., van Diest, P. J. & Pasterkamp, G. SlideToolkit: An Assistive Toolset for the Histological Quantification of Whole Slide Images. PLoS One 9, e110289 (2014).

11. Hamilton, P. W., Diest, P. J. Van, Williams, R. & Gallagher, A. G. Do we see what we think we see? The complexities of morphological assessment. 285–291 (2009). doi:10.1002/path

12. Polley, M.-Y. C. et al. An international Ki67 reproducibility study. J. Natl. Cancer Inst. 105, 1897–906 (2013).

13. Ruifrok, A. C. & Johnston, D. A. Quantification of histochemical staining by color deconvolution. Anal Quant Cytol Histol. 23, 291–9 (2001).

14. Galon, J. et al. Type, density, and location of immune cells within human colorectal tumors predict clinical outcome. Science 313, 1960–4 (2006).

15. Breiman, L. Random forests. Mach. Learn. 45, 5–32 (2001).

16. Munro, A. J., Lain, S. & Lane, D. P. P53 abnormalities and outcomes in colorectal cancer: a systematic review. Br. J. Cancer 92, 434–44 (2005).

17. McCluggage, W. G., Soslow, R. A. & Gilks, C. B. Patterns of p53 immunoreactivity in endometrial carcinomas: ‘all or nothing’ staining is of importance. Histopathology 59, 786–788 (2011).

18. Boyle, D. P. et al. The prognostic significance of the aberrant extremes of p53 immunophenotypes in breast cancer. Histopathology 1–13 (2014). doi:10.1111/his.12398

19. Garon, E. B. et al. Pembrolizumab for the treatment of non-small-cell lung cancer. N. Engl. J. Med. 372, 2018–28 (2015).

20. Pyo, J.-S., Kang, G. & Kim, J. Y. Prognostic role of PD-L1 in malignant solid tumors: a meta-analysis. Int. J. Biol. Markers 0 (2016). doi:10.5301/JBM.2016.16048

21. Dunne, P. D. et al. Immune-derived PD-L1 gene expression defines a subgroup of stage II/III colorectal cancer patients with favorable prognosis that may be harmed by adjuvant chemotherapy. Cancer Immunol. Res. 1–11 (2016). doi:10.1158/2326-6066.CIR-15-0302

22. Lee, L. H. et al. Patterns and prognostic relevance of PD-1 and PD-L1 expression in colorectal carcinoma. Mod. Pathol. 29, 1433–1442 (2016).

23. Huijbers, A. et al. The proportion of tumor-stroma as a strong prognosticator for stage II and III colon cancer patients: Validation in the victor trial. Ann. Oncol. 24, 179–185 (2013).

24. Mesker, W. E. et al. The carcinoma-stromal ratio of colon carcinoma is an independent factor for survival compared to lymph node status and tumor stage. Cell Oncol 29, 387–398 (2007).

25. Satyanarayanan, M., Goode, A., Gilbert, B., Harkes, J. & Jukic, D. OpenSlide: A vendor-neutral software foundation for digital pathology. J. Pathol. Inform. 4, 27 (2013).

26. R Core Team. R: A Language and Environment for Statistical Computing. (2016). at https://www.r-project.org/

27. Therneau, T. M. A Package for Survival Analysis in S. (2015). at https://cran.r-project.org/package=survival

28. Goulding, H. et al. A new immunohistochemical antibody for the assessment of estrogen receptor status on routine formalin-fixed tissue samples. Hum. Pathol. 26, 291–294 (1995).

29. Achanta, R. et al. SLIC Superpixels Compared to State-of-the-Art Superpixel Methods. Pattern Anal. Mach. Intell. IEEE Trans. 34, 2274–2282 (2011).

30. Haralick, R. M., Shanmugam, K. & Dinstein, I. Textural Features for Image Classification. IEEE Trans. Syst. Man. Cybern. (1973). doi:10.1109/TSMC.1973.4309314

